# Neurovegetative symptom subtypes in young people with major depressive disorder and their structural brain correlates

**DOI:** 10.1101/755447

**Authors:** Yara Toenders, Lianne Schmaal, Ben J. Harrison, Richard Dinga, Michael Berk, Christopher G. Davey

## Abstract

**Background:** Depression is a leading cause of burden of disease among young people. Current treatments are not uniformly effective, in part due to the heterogeneous nature of major depressive disorder (MDD). Refining MDD into more homogeneous subtypes is an important step towards identifying underlying pathophysiological mechanisms and improving treatment of young people. In adults, symptom-based subtypes of depression identified using data-driven methods mainly differed in patterns of neurovegetative symptoms (sleep and appetite/weight). These subtypes have been associated with differential biological mechanisms, including immuno-metabolic markers, genetics and brain alterations (mainly in the ventral striatum and insular cortex).

**Methods:** K-means clustering was applied to individual depressive symptoms from the Quick Inventory of Depressive Symptoms (QIDS) in 275 young people (15-25 years old) with MDD to identify symptom-based subtypes, and in 244 young people from an independent dataset (a subsample of the STAR*D dataset). Insula surface area and thickness and ventral striatum volume were compared between the subtypes using structural MRI.

**Results:** Three subtypes were identified in the discovery dataset and replicated in the independent dataset; severe depression with increased appetite, severe depression with decreased appetite and severe insomnia, and moderate depression. The severe increased appetite subtype showed lower surface area in the anterior insula compared to both healthy controls and the moderate subtype.

**Conclusions:** Our findings in young people replicate the previously identified symptom-based depression subtypes in adults. The structural alterations of the anterior insular cortex add to the existing evidence of different pathophysiological mechanisms involved in this subtype.

## Introduction

Approximately 322 million people worldwide (5% of the world’s population) suffer from Major Depressive Disorder (MDD), a disease characterized by a depressed mood and associated symptoms (1). In young people, depressive disorders are the main cause of global burden of disease (2). The onset of MDD peaks during adolescence and young adulthood, and earlier onset of MDD is associated with decreased quality of life and increased impairment in social and occupational functioning later in life (3, 4). Currently available treatments are not uniformly effective for adolescent depression, with response rates around 61% for antidepressants and 55% for psychotherapy (5, 6). The unpredictable nature of treatment response might be explained, at least in part, by the heterogeneity of MDD.

The most commonly used systems for classifying mental disorders, The ICD 10 (International Classification of Diseases 10th revision) and Diagnostic and Statistical Manual of Mental Disorders (DSM-5), categorize a broad spectrum of depressive symptom patterns within a single MDD diagnosis. To receive an MDD diagnosis, a minimum of 5 of the 9 DSM criteria for MDD have to be met. Considering that some of the criteria include symptoms of opposite polarity (e.g. increased versus decreased appetite, weight gain versus loss, insomnia versus hypersomnia, and psychomotor agitation versus retardation), almost 1500 different combinations of MDD symptoms lead to the same DSM diagnosis of MDD (7). Thus, patients with the same diagnosis show heterogeneous depressive symptom profiles, which may reflect different underlying neurobiological mechanisms that could require different treatments. As people with different phenotypic presentations of depression are not uniquely identified and diagnosed, they cannot be stratified into different treatments.

Several attempts to identify subtypes of depression have been made to overcome these issues associated with the traditional diagnostic classification and the heterogeneity of MDD. Traditionally, subtypes of depression have been defined based on subjective expert consensus. An example of describing different subtypes are the DSM atypical and melancholic depression specifiers (1). The atypical specifier is characterized by mood reactivity in combination with increases in weight or appetite, hypersomnia and/or leaden paralysis. The melancholic specifier is distinguished by opposite neurovegetative symptoms: decreases in weight or appetite and early morning awakening, in addition to psychomotor agitation or retardation, worse mood in the morning and excessive feelings of guilt. Existence of the melancholic specifier has been confirmed by prior research, however, the atypical specifier has been questioned. For example, mood-reactivity, the only obligatory atypical symptom in the DSM, does not show associations with the other atypical features (8, 9).

More recently, data-driven approaches have been employed to identify symptom-based depression subtypes. Replicated across a number of studies in adults, latent class analysis has derived data-driven typical versus atypical neurovegetative symptom subtypes (10–17). The typical and atypical subtypes are usually characterized as having similar affective and cognitive symptoms, differing only on sleep and appetite profiles. Of note, the atypical neurovegetative symptom subtype differs from the atypical specifier in the DSM as it mainly shows only reversed neurovegetative symptoms (i.e., increased appetite and weight, and in some studies also hypersomnia). However, not much is known about whether similar subtypes exist in young people with depression, although one study suggests they are similar to adults (18).

Importantly, there are biological differences between the data-driven neurovegetative symptom subtypes in adults. Higher levels of leptin, inflammatory markers (C-reactive protein (CRP), interleukin-6 (IL-6), interleukin 1 receptor antagonist (IL-1RA) and tumor necrosis factor-α (TNF-α)), insulin and higher BMI are associated with the atypical or increased appetite subtype (15, 19–21). Conversely, higher cortisol and ghrelin levels are associated with the more typical subtype characterized by decreased appetite (19, 21–23). In addition, genetic studies have shown that the atypical subtype with increased appetite is associated with a higher polygenic risk score for BMI, leptin and CRP, whereas the typical subtype showed a stronger association with polygenic risk scores for psychiatric disorders such as schizophrenia (24, 25). Moreover, brain activation responses to pictures of food have been shown to differentiate depressed patients selected on having either increased or decreased appetite (21, 26). Higher cortisol levels in the MDD group with decreased appetite were negatively correlated with ventral striatal activity during a food task; while the MDD group with increased appetite showed a positive correlation between insulin resistance and posterior and dorsal mid-insula cortex activity (21). In addition, higher anterior insula cortex activity in response to these appetitive food pictures was observed in the MDD group with increased appetite (26).

There is now consistent evidence in adults for different biological correlates across neurovegetative symptom subtypes, however, it remains unknown whether similar subtypes exist in young people and if they are characterized by similar biological mechanisms. The current study aims to replicate the data-driven symptom profiles based on neurovegetative symptoms, previously identified in adults, in young people with MDD. Furthermore, we included an additional independent sample as a replication cohort. In addition, the study aims to examine structural brain alterations associated with the identified subtypes. We hypothesize that similar subtypes exist in young people, mainly distinguished by opposite neurovegetative symptoms. We also hypothesize that structural alterations in subregions of the insula and ventral striatum, regions that have been implicated in subtype differences in previous adult studies (21, 26), may differentiate between the symptom subtypes.

## Methods and Materials

### Participants

#### Discovery sample

Participants were recruited from youth mental health centers in Australia as part of the YoDA-A and YoDA-C (Youth Depression Alleviation) studies (27, 28). All participants were aged between 15 and 25 years old and diagnosed with a primary diagnosis of MDD. In total, 275 young people with MDD and 100 age and gender matched healthy controls (HC) were included. The HC participants were recruited through advertisements and did not have a present or past diagnosis of MDD or anxiety disorders. Participants were excluded if they suffered from an acute medical disorder, had experienced any psychotic episodes, or were diagnosed with bipolar disorder. Further, pregnancy, breastfeeding and any contraindications to MRI were exclusion criteria. The participants gave written informed consent and the Melbourne Health Human Research Ethics Committee approved the study protocol.

#### Replication sample

The data for the independent replication sample came from the STAR*D study, a large multicenter study examining antidepressant effectiveness (29). To match the YoDA sample, the sample was restricted to young people between 18 and 25 years old with an MDD diagnosis, resulting in a sample size of 244. All participants scored 14 or higher on the Hamilton Depression Rating Scale (HDRS) (30), indicating moderate to severe depression. Exclusion criteria were a primary diagnosis of schizophrenia, bipolar disorder, anorexia nervosa, bulimia or obsessive-compulsive disorder. In addition, the participants were free of antidepressants when they entered the study.

### Procedure

#### Discovery sample

The YoDA participants were screened using the Structured Clinical Interview for the DSM-IV (SCID) during the baseline assessment (29). In addition, depressive symptoms were measured using the Montgomery–Åsberg Depression Rating Scale (MADRS) as well as the Quick Inventory of Depressive Symptomatology Self Report (QIDS-SR) (31, 32). A score of 20 or higher on the MADRS, indicating at least moderate severity of symptoms, was an inclusion criterion in these studies. Further, participants completed the Generalized Anxiety Disorder 7 questionnaire (GAD-7), Social and Occupational Functioning Assessment Scale (SOFAS), Alcohol Use Disorder Identification Test (AUDIT) (33–35) and other questionnaires not germane to this study. Within two weeks of the baseline assessment, and prior to commencing the study treatments, a subset of the participants (137 MDD patients and 100 healthy controls) underwent a structural MRI scan.

#### Replication sample

Baseline scores of the QIDS of STAR*D participants were included in this study (31). The Psychiatric Diagnostic Screening Questionnaire (PDSQ) was used as a diagnostic screening tool in the STAR*D study (36). Quality of life was assessed with the Quality of Life Enjoyment and Satisfaction Questionnaire (Q-LES-Q) and impaired functioning using the Work and Social Adjustment Scale (WSAS) (37, 38). No MRI data was available in this sample.

### MRI data

#### Discovery sample

The T1 weighted scan lasted ∼4 minutes and was performed on a 3T General Electric Signa Excite at Sunshine Hospital (Western Health, Melbourne). An 8-channel phased-array head coil was used (TR: 7900 ms, echo time: 3000 ms, thickness (no gap): 1 mm, flip angle: 13°, field of view: 25.6 cm, matrix: 256×256 pixels).

The cortical parcellation was performed using FreeSurfer (version 5.3) (39). The segmentations and parcellations were visually inspected and outliers were examined using the ENIGMA protocol (http://enigma.ini.usc.edu/protocols/imaging-protocols). Cortical surface area and cortical thickness of the anterior and posterior insula (based on the Destrieux atlas, (40)) and ventral striatal (nucleus accumbens) volume in the left and right hemisphere were included as regions of interest.

### Data analysis

#### Symptom subtypes

A k-means clustering in R was applied to the 16 depression items of the QIDS to identify symptom subtypes in the young people diagnosed with MDD from the YoDA sample (41). With k-means clustering, clusters are formed based on the cluster mean (centroid) that is closest to a data point (the item score of a subject) to keep the centroids as small as possible. The QIDS item data were scaled to make sure all variables had the same weight.

##### Selecting the optimal number of clusters

The number of clusters (k) was selected based on the highest number of partitioning methods in R that selected the same optimal number of clusters. The R package NbClust was used to determine the optimal number of clusters by looking at different combinations of distance measures and clustering methods based on hierarchical clustering (ward.D2, which minimizes variance within clusters) (for more details, please see supplemental material) (42).

##### K-means clustering

The optimal number of clusters determined by NbClust was used as an input parameter to K-means clustering using the stats package in R (41), to identify the centers of the clusters (centroids). To prevent the clustering settling on local minima, an initialization method was used to pick cluster means that covered the full range (43, 44). In this initialization method, random centers were selected, after which the procedure was reran to readjust the centers. The centroids of the next cluster were selected by maximizing the distance to the centroids that were selected before. These centers were used to run the K-mean clustering.

##### Testing validity and stability

Three methods were used to test the validity and reliability of the clusters. First, to test the stability of the clusters we repeated our clustering analysis in 10,000 randomly selected subsamples, each containing 100 participants from a pre-selected training sample (which consisted of 70% of the total sample). In each of the 10,000 subsamples, participants left out of the cluster identification process (the remaining 30%) were assigned to clusters using linear discriminant analysis classifiers. The left-out sample was combined with the training sample to form a complete cluster solution. We then tested whether the individual cluster assignments were stable over the 10,000 subsamples by calculating an adjusted Rand score to test the similarity between each subsampling clustering solution compared to the original clustering solution. A Rand index of 1 means that the clustering solutions completely agree on the labels, while a Rand-index of 0 represents a disagreement in the clustering. We also calculated the cluster-to-cluster index, which represents the mean distance between the clusters in the original and the new clustering obtained through resampling. Second, the optimal number of clusters was tested against a null distribution with permutation testing (45). The same analysis procedure, including subsampling and permutation testing was repeated in the independent replication dataset STAR*D. A latent class analysis was performed to test the robustness of the k-means clustering method and to compare our findings to findings in previous adult studies that mainly employed latent class analysis (Supplementary Materials).

#### Differences in clinical and structural brain characteristics between subtypes

The symptom subtypes that were identified were compared on clinical and demographic characteristics in R using ANOVA or chi-square tests, and if there was a significant difference at *p*<0.05, *posthoc* tests with Tukey HSD correction for multiple comparisons (3 tests to compare the 3 subtypes).

In the YoDA sample, anterior and posterior insula cortex surface area and thickness and ventral striatum volume (in the left and right hemisphere) were compared between subtypes using an ANCOVA with group (MDD subtypes and healthy controls) as predictor and age and sex, as well as intercranial volume (ICV) as covariates. We did not include ICV as a covariate in analysis with insula thickness measures, since thickness does not scale with head size (46). False discovery rate (FDR) correction was applied and Tukey HSD corrected *posthoc* tests were performed when a significant main effect of group was found. Analyses were repeated in an antidepressant naïve sample to control for the possible effect of antidepressant use.

## Results

### Symptom subtypes

A 3-cluster solution was found to be the optimal fit according to 9 out of 26 partitioning methods (see Supplemental Figure S1). The stability of the clusters was tested for 2 as well as 3 clusters. The average Rand Index was 0.40 for 2 clusters, and 0.55 for 3 clusters. In addition, the average cluster-to-cluster distance was 1.35 for 2 clusters and 1.85 for 3 clusters, meaning that more labels agreed when 3 clusters were used and the means of different clusters were further apart in the 3-cluster solution. In addition, the Scott and Friedman partitioning measure showed that the index number for this number of clusters was higher than the cluster indices of an empirical null distribution, meaning that 3 clusters described the data better than data with no underlying clusters (Scott: p<0.001, Friedman: p=0.01, Supplemental Figure S2).

The clusters were labelled as following: moderate depression (n=111, MOD), severe depression with increased appetite (n=59, SIA) and severe depression with decreased appetite and insomnia (n=105, SDA) (Figure 1A). The MOD subtype endorsed symptoms such as a sad mood, lack of general interest, fatigue and typical neurovegetative symptoms of decreased appetite, weight loss and insomnia. The SIA and SDA subtypes both showed a higher severity of symptoms overall than the MOD subtype. The SIA subtype was uniquely characterized by endorsement of reversed (atypical) neurovegetative symptoms of increased appetite and weight gain, whereas the SDA subtype showed decreased appetite and higher levels of insomnia. The SIA subtype consisted of more females and was associated with higher BMI compared to SDA and MOD (Table 1). Similar clusters were identified in the STAR*D dataset, including a moderate depression subtype (MOD, n=108), a severe depression with increased appetite and weight gain subtype (SIA, n=54) and a severe depression with decreased appetite and highest levels of insomnia subtype (SDA, n=82) (Figure 1B, Table 2).

**Table 1.**
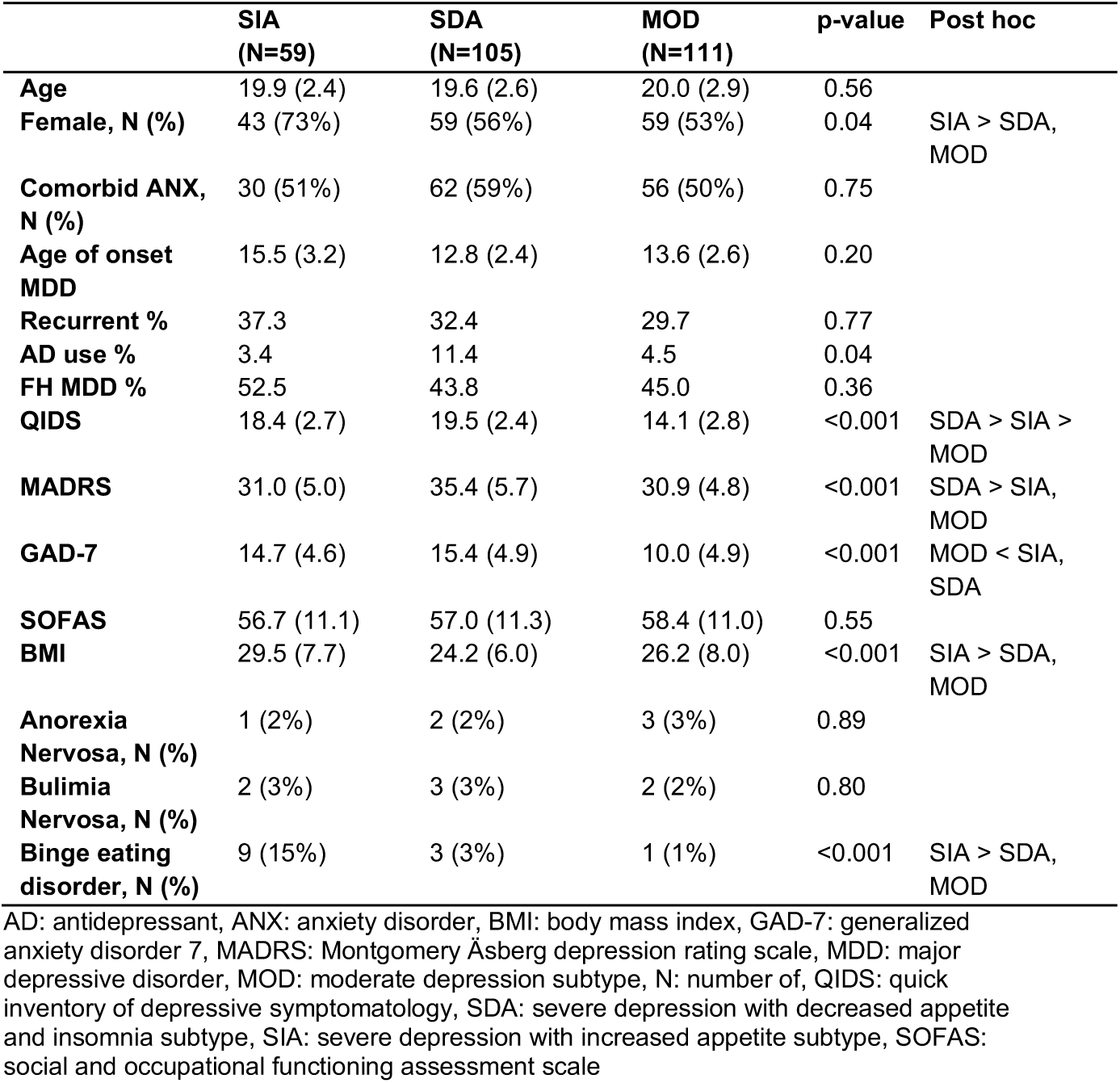
Demographics and clinical characteristics of symptom subtypes identified in the YoDA discovery sample.

**Table 2.** Demographics and clinical characteristics of symptom subtypes identified in the STAR*D replication sample.

|  | **SIA**<br>**(N= 54)** | **SDA**<br>**(N= 82)** | **MOD**<br>**(N=108)** | **p-value** | **Post hoc** |
| --- | --- | --- | --- | --- | --- |
| **Age** | 22.3 (2.0) | 21.8 (2.0) | 21.8 (1.9) | 0.29 |  |
| **Female, N (%)** | 42 (78%) | 60 (73%) | 75 (69%) | 0.53 |  |
| **Comorbid ANX, N (%)** | 5 (9%) | 11 (13%) | 16 (14%) | 0.61 |  |
| **Eating disorder, N (%)** | 1 (2%) | 1 (1%) | 0 (0%) | 0.41 |  |
| **QIDS** | 18.3 (3.2) | 18.4 (2.5) | 12.4 (2.3) | <0.001 | SIA, SDA > MOD |
| **Hamilton** | 21.1 (4.4) | 24.4 (5.2) | 19.8 (4.0) | <0.001 | SDA > SIA, MOD |
| **Q-LES-Q** | 38.8 (10.7) | 38.6 (12.6) | 50.2 (13.4) | <0.001 | SIA, SDA < MOD |
| **WSAS** | 26.0 (7.4) | 25.3 (7.1) | 17.5 (7.2) | <0.001 | SIA, SDA > MOD |

ANX: anxiety disorder, MOD: moderate depression subtype, N: number of, QIDS: quick inventory of depressive symptomatology, Q-LES-Q: quality of life enjoyment and satisfaction questionnaire, SDA: severe depression with decreased appetite and insomnia subtype, SIA: severe depression with increased appetite subtype, WSAS: work and social adjustment scale

**Figure 1.**
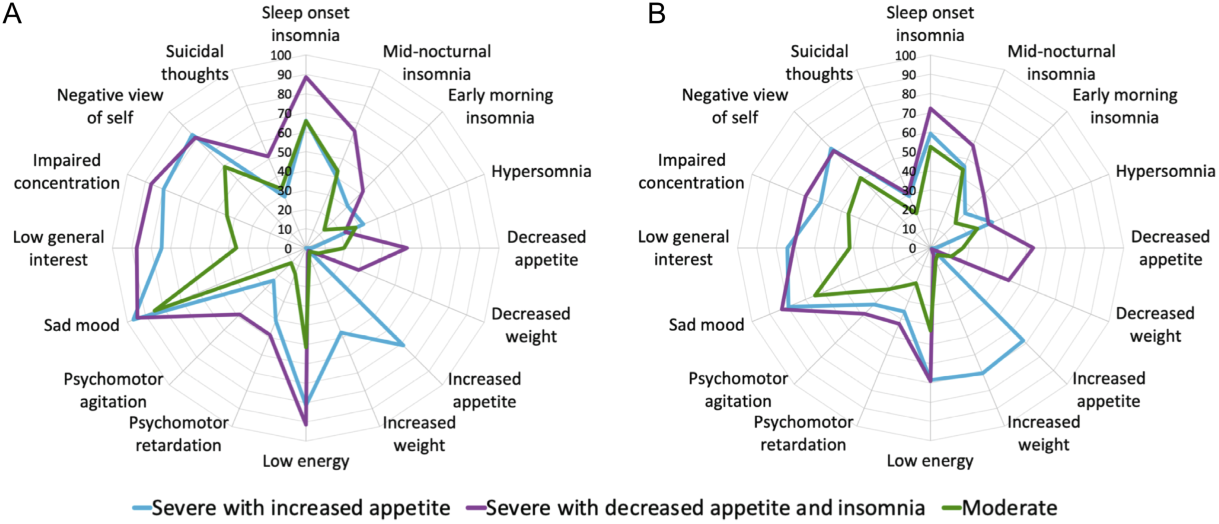
Symptom subtypes in the YoDA discovery sample (A) and in a subsample of the STAR*D replication sample (B). A severe depression with increased appetite (SIA) subtype, severe depression with decreased appetite and insomnia (SDA) subtype and a moderate depression (MOD) subtype were identified in both datasets. The axis shows the percentage of subjects within a subtype that shows the symptoms in the radar plot (QIDS items).

AD: antidepressant, ANX: anxiety disorder, BMI: body mass index, GAD-7: generalized anxiety disorder 7, MADRS: Montgomery Äsberg depression rating scale, MDD: major depressive disorder, MOD: moderate depression subtype, N: number of, QIDS: quick inventory of depressive symptomatology, SDA: severe depression with decreased appetite and insomnia subtype, SIA: severe depression with increased appetite subtype, SOFAS: social and occupational functioning assessment scale

### Neurobiological alterations in symptom subtypes

Left and right anterior insula surface area showed a main effect of group (p=0.01), which was driven by lower surface area in these regions in the SIA subtype compared to healthy controls (*posthoc*: left anterior insula: p=0.05, right anterior insula: p=0.03) (Figure 2).

**Figure 2.**
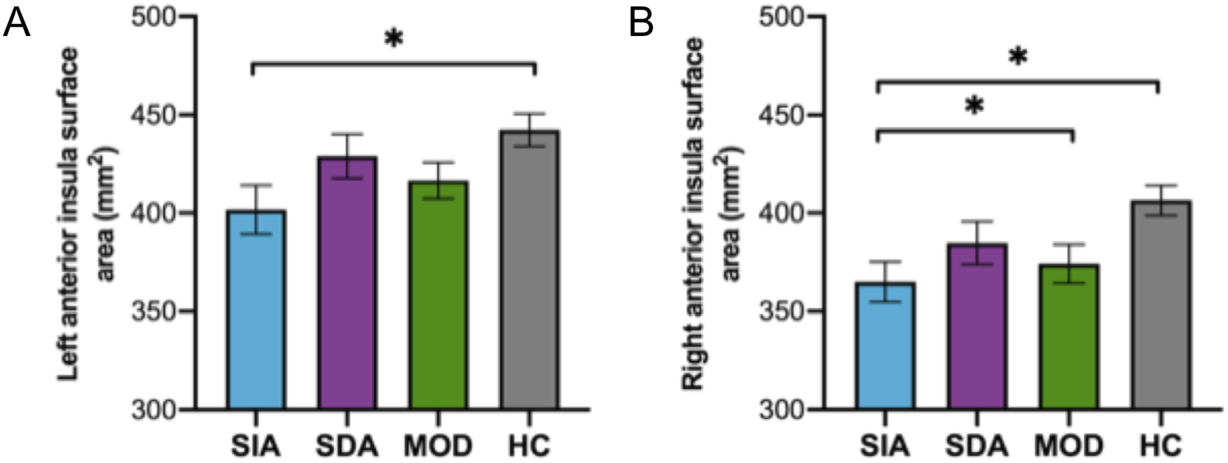
Left and right anterior insula surface area in the subtypes in the YoDA discovery sample. Severe increased appetite (SIA) subtype, Severe decreased appetite and insomnia (SDA) subtype, Moderate (MOD) subtype and Healthy control (HC). SIA showed significantly lower surface area in the left anterior insula compared to HC. In the right anterior insula, SIA showed lower surface area compared to HC and MOD.

Additionally, right anterior insula surface area was also lower in the SIA subtype compared to the MOD subtype (marginally significant at p=0.05). No differences were found in posterior insula surface area, thickness of insular subregions and ventral striatum volume. Among the participants with neuroimaging data, the subtypes did not show differences in BMI (see Supplemental Table S1). The results were replicated in a subset of the sample excluding lifetime antidepressant users. In the subset consisting of only antidepressant naïve patients (n=97), the main effects for left and right anterior insula were still significant (p=0.03 and p=0.03). However, the SIA subtype showed less robust differences from healthy controls (p=0.06 and p=0.08) and the surface area in the left and right anterior insula in the SIA subtype does not differ from the moderate subtype in this sample (p=0.27 and p=0.17).

## Discussion

The aims of the current study were to identify data-driven subtypes of depressive symptoms in young people (aged 15-25) with MDD and to compare potential structural brain alterations between these subtypes. The data-driven symptom subtypes found in the YoDA study cohort were in line with the subtypes characterized by opposite neurovegetative symptoms previously identified in adults (11–18). One subtype showed atypical or reversed neurovegetative symptoms, mainly discriminated by increased appetite and weight gain, and two subtypes showed typical neurovegetative symptoms, including insomnia, decreased appetite and weight loss, with the typical symptom subtypes having different levels of overall severity (moderate versus severe). We replicated these data driven symptom subtypes in a subsample of the STAR*D study, an independent sample of MDD patients within a similar age range. Symptom-based subtypes in young people have only been studied in one previous study in adolescents, that used latent class analysis to identify similar subtypes (18).

The data used in the current study is unique, since 34% of the MDD patients with imaging data were diagnosed with their first episode of MDD and 70% were antidepressant naïve. Identifying similar subtypes as found in adults in these more clinically specific (antidepressant free and at an early stage of the disorder) young people further validates the existence of the subtypes. Moreover, identifying similar subtypes in adolescents and young adults is relevant, since mood, appetite and sleep and their underlying biological processes go through developmental changes in adolescence, such as maturation of neural emotion regulation and reward processing circuitries, increasing levels of leptin, and a shift in the circadian rhythm (47–49).

In line with previous adult studies, the increased appetite subtype had more females and a higher BMI than the other subtypes (11, 13, 16, 18). In addition, unlike some adult studies, the increased appetite subtype we identified was not discriminated by hypersomnia. However, hypersomnia items in a self-report questionnaire show low correlations with objective sleep measures (50, 51). Additionally, whereas three items assess insomnia in the QIDS self-report, only one item targets hypersomnia, and the disturbances might be more complex than assessed in that single question (for example fractionated or irregular night-time sleep but increased duration of sleep, including daytime napping). Therefore, sleep disturbances may still exist in the subtype with atypical neurovegetative symptoms. More ecologically valid assessments of sleep disturbances should be employed in future studies to examine sleep disturbances in these subtypes.

This study is the first to examine differences in structural brain alterations between data-driven symptom-based subtypes. We found lower anterior insula surface area in the increased appetite subtype compared to healthy controls and the moderate severity subtype. Different parts of the insula are thought to have different roles, with the anterior insula important for integration of interoceptive information and reward and motivational processes (52). The anterior portion of the insula is preferentially interconnected with the orbitofrontal (OFC) and anterior cingulate cortices (ACC) and ventral striatum (53–55). Together with the dorsal ACC, the anterior insula forms a core hub of the so-called ‘salience network’, commonly implicated in interoceptive awareness; integrating external and internal stimuli to guide an individual’s actions and decisions (56–59). The anterior insula integrates information about the motility of the digestive system and hunger. Hormones (including leptin and insulin), body weight status, and inflammation have been shown to influence insula activity and volume (60–63). Since alterations in hormones, such as leptin and insulin, and inflammatory markers have been observed in the increased appetite subtype in adults and those hormones and inflammatory markers have been found to affect surface area (64), is possible that alterations in these endocrine factors that may be unique to this atypical neurovegetative subtype affect surface area.

Furthermore, the insula is implicated in reward processing and emotion regulation, processes that have been associated with food intake (65–68). Previous research reported increased brain activity in the anterior insula and other reward regions including the ventral striatum in response to pictures of food in adult MDD patients with increased appetite (26). In addition, emotion regulation disturbances have shown to increase emotional eating (69), which may underlie the increased appetite and weight gain observed in the atypical neurovegetative subtype, potentially mediated by structural alterations in the anterior insula (70).

Only two prior studies in adults examined differences in brain measures between MDD patients selected on the presence of depression-related symptoms of increased appetite versus decreased appetite (21, 26). These studies examined neural responses during an fMRI food picture task, and found that lower ventral striatum activity was associated with higher cortisol in the decreased appetite subtype. In contrast, in the increased appetite subtype higher anterior insula activity was observed. In line with these studies by Simmons et al., we found anterior insula surface area alterations in the increased appetite subtype. However, no differences in ventral striatum volume between the subtypes were found in the present study, suggesting that alterations in the ventral striatum might be restricted to a functional level.

Interestingly, only surface area differences were observed for the anterior insula and no differences were found in insular cortical thickness. Cortical surface area and cortical thickness are two distinct characteristics of the brain’s cortex and have different developmental pathways. Cortical thickness increases until approximately age 2, whereas cortical surface area increases, depending on the region, until adolescence, making it more vulnerable to early life stressors (71–74). In addition, cortical surface area alterations have been found to be associated with early onset depression (75, 76), and prior research shows that the increased appetite, or atypical neurovegetative, subtype is associated with earlier onset of depression (13, 19, 24). However, since this study consisted of adolescents and young adults, the age of onset was low overall and did not differ between subtypes.

A few limitations of the study should be noted. The exclusion criteria of the YoDA studies might have influenced clustering results, and compromise generalizability. Only young people with MDD who showed moderate to severe depressive symptoms were included, therefore not representing the whole depressive spectrum. Additionally, the k-means clustering might have been affected by the high negative correlations between the increased appetite/weight and decreased appetite/weight symptoms, and between insomnia and hypersomnia. There has been some critique regarding the subtyping based on symptoms including these opposite neurovegetative symptoms using a latent class analysis or other data-driven techniques, since they are complete opposites and one symptom automatically rules out the possibility of showing the other symptom (e.g., a person can’t endorse both weight gain and weight loss at the same time, although they can endorse no changes in weight). The negative correlations between increased and decreased appetite (−0.49) and weight (−0.31), and between insomnia and hypersomnia (−0.14 to −0.21), known as a violation of conditional independence, have likely dominated the clustering and may have masked subtypes based on other patterns of symptom endorsement (77). However, differences in genetics, blood markers of inflammation, leptin insensitivity and insulin resistance, and neuroimaging markers have been repeatedly found between these subtypes derived using a data-driven analysis (19, 20, 24, 25, 78 and the current study) and when selected on the presence of increased versus decreased appetite (21, 26). Therefore, the subtypes seem clinically relevant.

This clinical relevance of these neurovegetative symptom subtypes is underlined by the persistence of sleep and appetite disturbances after treatment for depression (79, 80). In addition, these neurovegetative symptoms do not affect clinical management decisions as much as mood symptoms, even though they are associated with high risks of suicide (81), obesity and metabolic syndrome (82), and depression recurrence (83–85). Subtyping depression based on neurovegetative symptoms could lead to more targeted intervention. However, to achieve a more personalized intervention, future research should investigate whether the different subtypes based on neurovegetative symptoms respond differently to traditional (e.g. psychotherapy, antidepressants) and novel treatments. Our findings of structural alterations in the anterior insula together with previous findings of functional alterations in the insula uniquely associated with the atypical neurovegetative subtype suggest that core functions of the insula including interoceptive function, emotion regulation and reward processing may be promising treatment targets for this specific subtype of depression.

To conclude, we were able to replicate the existence of reversed neurovegetative and typical neurovegetative symptom subtypes of depression in two adolescent/young adult MDD samples. This was the first study to show that these symptom subtypes were associated with cortical surface alterations in the anterior insula, with the increased appetite showing lower surface area compared to the moderate subtype and healthy controls. Together with previous findings in adults, our current findings suggest that the subtype with atypical neurovegetative symptoms may have a unique biological signature. Moreover, neurovegetative symptoms are associated with poorer clinical outcomes and antidepressant treatment have shown to work more effectively for mood and cognitive symptoms than for atypical and sleep symptoms (86, 87). Therefore, these neurovegetative symptoms subtypes, characterized by changes in sleep and appetite, should be noted when treating young people with depression.

## Acknowledgements

This study was supported by National Health and Medical Research Council of Australia (NHMRC) Project Grants to CGD (YoDA C ID 1024570), MB (YoDA A ID 1027315) and BJH (1064643). BJH is supported by a NHMRC Career Development Fellowship (1124472). supported by an NHMRC Career Development Award (141738). MB is supported by a National Health and Medical Research Council (NHMRC) Senior Principal Research Fellowship (APP1059660 and APP1156072). LS is supported by a NHMRC Career Development Fellowship (1140764) and gratefully acknowledges support from The Netherlands Brain Foundation (F2014[1]-24) for this work.

