## Supplemental Material for "Neurovegetative symptom subtypes in young people with major depressive disorder and their structural brain correlates"

**Supplementary materials**

**Methods**

*Optimal number of clusters compared to a null distribution using permutation testing*

The partitioning methods (such as Scott, Friedman or Dunn) from the NbClust package in R that choose a certain number of clusters, do not test whether this number of clusters is significantly better at explaining the data than it would be under the null hypothesis of data without any underlying clusters. Therefore the statistical significance of the index of 3 partition methods were tested (1, 2). In this procedure, the null hypothesis is that the data came from a single 2-dimensional Gaussian distribution (i.e. distribution with no underlying clusters). Second, we repeatedly took 10,000 random samples from a bivariate Gaussian distribution defined by a covariance matrix of the QIDS (Quick Inventory of Depressive Symptomatology) items. Third, we ran the same hierarchical clustering procedure as we performed on the real data on each random sample and calculated the best obtained partitioning indices, thus obtaining an empirical null distribution of these indices. The p-value was then defined as the proportion of the calculated indices in the null distribution smaller than what we observed in the real data.

*Latent class analysis*

To test how robust the k-means clustering method was, a latent class analysis (LCA) using the ‘poLCA’ package in R, was performed on the YoDA and STAR*D data as well (3). The LCA clusters people into different classes, based on an underlying latent variable that explains associations among the items. The optimal number of clusters was selected based on the lowest Bayesian Information Criteria (BIC).

**Results**

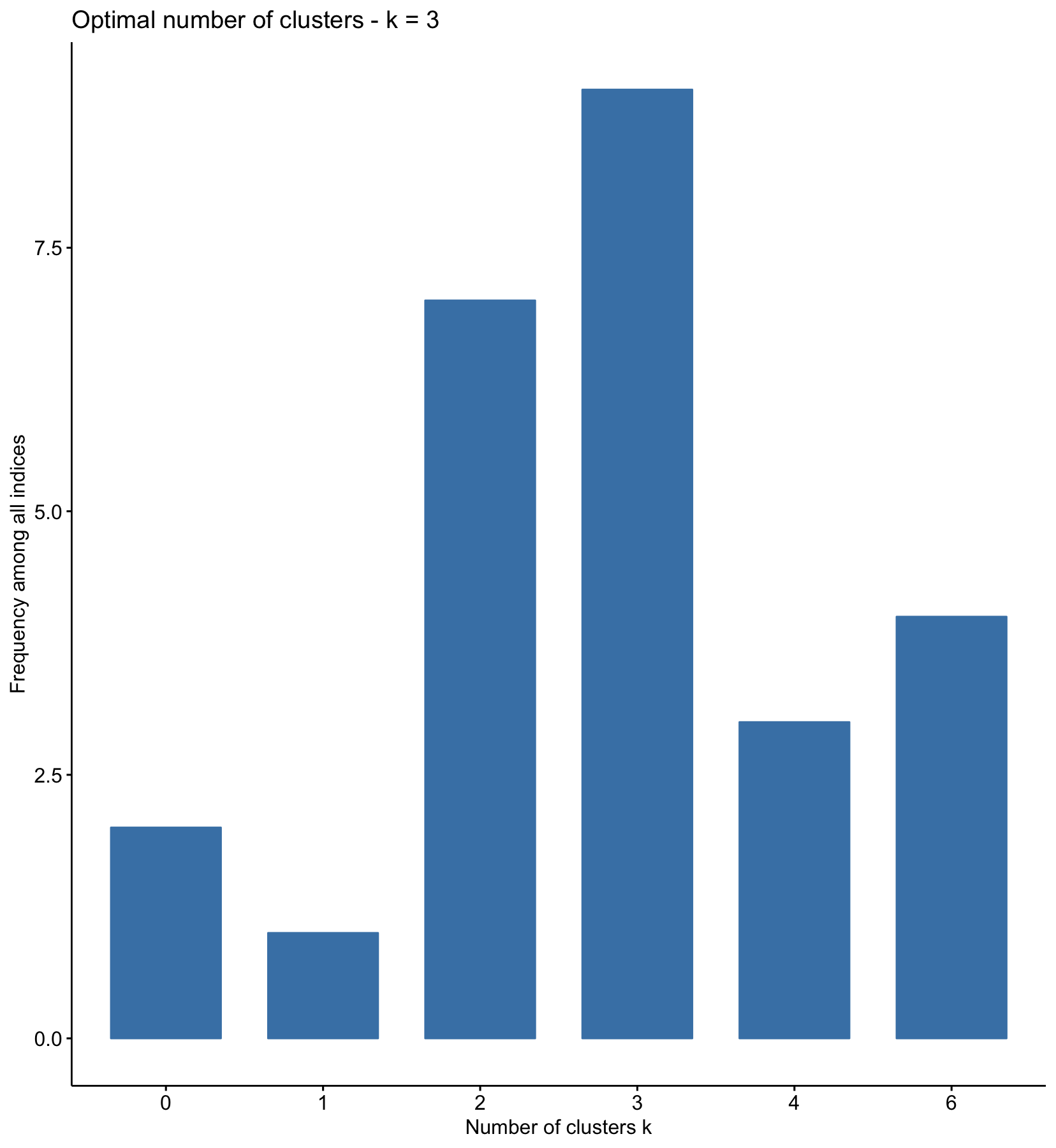

**Supplemental Figure S1. Optimal number of clusters in YoDA sample**

A 3-cluster solution was found to be the optimal fit according to 9 out of 26 partitioning methods (Supplemental Figure S1). A 2-cluster solution was also found to be the optimal fit according to 7 partitioning methods. Using permutation tests, the significance of the number of clusters in a null distribution was determined. For example, the Dunn, Scott and Friedman method suggested 3 clusters was the optimal solution, and 2 of them showed this was significant (p=0.990, p=0.001, p=0.009, Supplemental Figure S2). Conversely, CH, Silhouette and Pseudot2 showed that 2 clusters was the optimal solution, but these were not significantly different than the same indices within a null distribution (p=0.445, p=0.300, p=0.213).

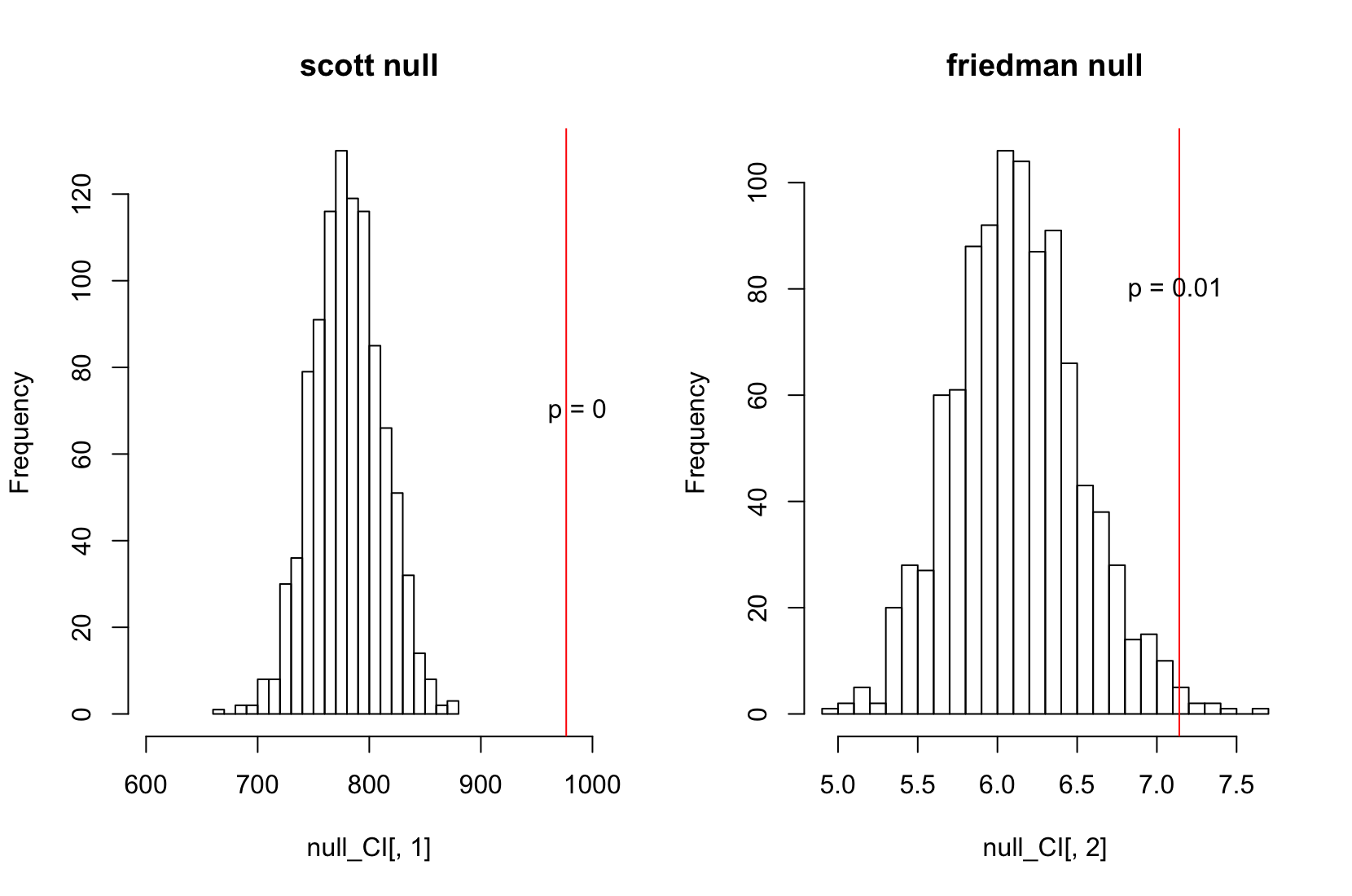

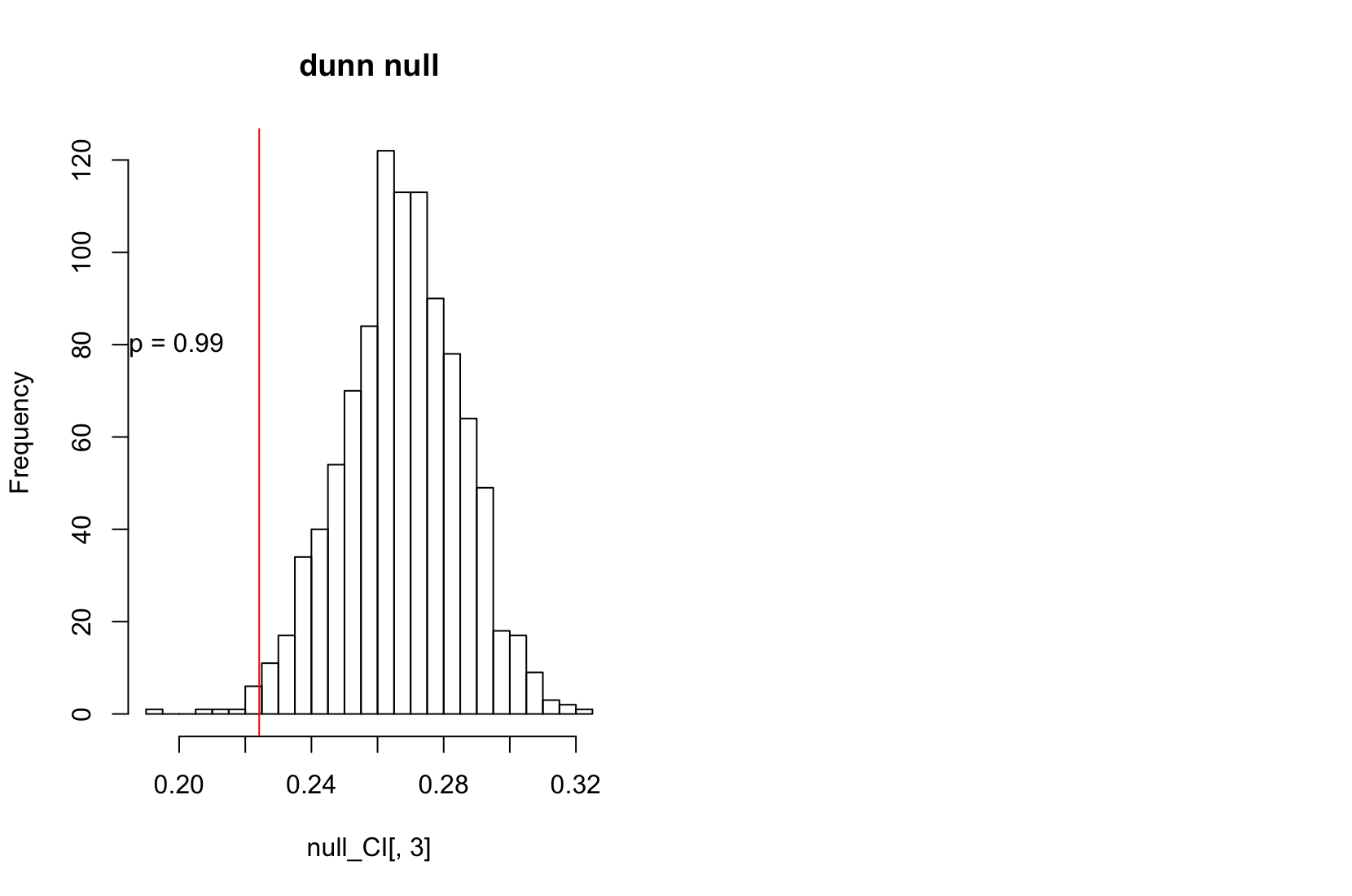

**Supplemental Figure S2. Significance of number of clusters in YoDA (3 clusters).**

*Symptom subtypes STAR*D*

Out of the 26 partition methods determining the optimal number of clusters, 8 clusters selected 2 clusters as optimal and 7 clusters selected 3 clusters. The stability was higher with 3 clusters (Rand Index 0.60 and cluster to cluster distance 1.25 compared to 0.27 and 2.90 for 2 clusters). Of the three measures used before in the YoDA sample, two (Dunn and Friedman) again selected 3 clusters as the optimal solution. The permutation testing that this was only significant in Friedman (p=0.223, p=0.001).

*Latent class analysis*

The two or three class solution seemed to be the best fit in the YoDA sample based on the BIC (BIC = 4947.4 and 4952.5). Because 3 classes were in line with the rest of the results, the 3 classes were chosen as the final solution. The 3 classes found were similar to the 3 clusters in the k-means clustering, again including a moderate depression subtype (N = 116), a severe typical vegetative symptoms subtype (N = 93) and an atypical vegetative symptom subtype (N = 66) (see Supplemental Figure S3). This was replicated in the STAR*D dataset (SIA = 55, SDA = 80, MOD = 109) (Supplemental Figure S3).

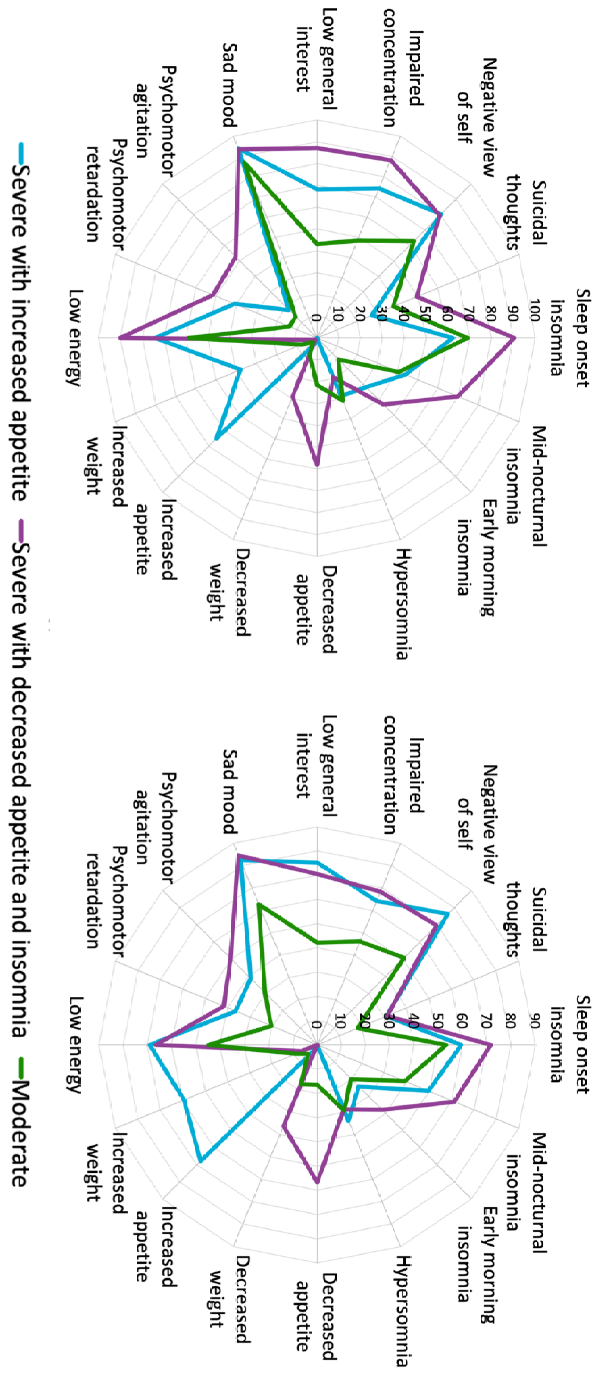

**Supplemental** **Figure S3. Symptom subtypes YoDA (left) and STAR*D (right) created with latent class analysis.** A severe depression with increased appetite (SIA) subtype, severe depression with decreased appetite (SDA) subtype and a moderate depression (MOD) subtype were identified in both datasets. The axis shows the percentage of subjects within a subtype that shows the symptoms in the radar plot (QIDS items).

**Supplemental Table S1. Demographics and clinical characteristics per symptom subtype in the subset of the YoDA discovery sample with imaging data.**

|  | SIA  (N=33) | SDA  (N=49) | MOD  (N=55) | p-value | Post hoc |
| --- | --- | --- | --- | --- | --- |
| Age | 19.7 (2.7) | 19.9 (2.6) | 19.5 (3.0) | 0.82 |  |
| Female, N (%) | 25 (75%) | 26 (53%) | 26 (47%) | 0.03 | SIA > SDA, MOD |
| Age of onset MDD | 15.4 (3.2) | 15.0 (2.5) | 15.8 (2.6) | 0.34 |  |
| MADRS | 31.0 (5.0) | 35.3 (5.7) | 31.6 (4.0) | <0.001 | SDA > SIA, MOD |
| QIDS | 18.5 (2.2) | 19.7 (2.5) | 14.0 (2.7) | <0.001 | SDA, SIA > MOD |
| BMI | 28.0 (7.3) | 24.8 (6.1) | 25.0 (6.5) | 0.10 |  |

BMI: body mass index, MADRS: Montgomery Äsberg depression rating scale, MDD: major depressive disorder, MOD: moderate depression subtype, N: number of, QIDS: quick inventory of depressive symptomatology, SDA: severe depression with decreased appetite and insomnia subtype, SIA: severe depression with increased appetite subtype
